# Characterization of mannitol metabolism genes in *Saccharina* explains its key role in mannitol biosynthesis and evolutionary significance in Laminariales

**DOI:** 10.1101/243402

**Authors:** Shan Chi, Tao Liu, Cui Liu, Yuemei Jin, Hongxin Yin, Xin Xu, Yue Li

## Abstract

As a unique photosynthetic product in brown algae, mannitol exhibits high synthesis and accumulation in *Saccharina japonica*. Mannitol acts as a carbon-storage compound and is an osmoprotectant, imparting increased tolerance to osmotic stress. However, the underlying biochemical and molecular mechanisms in macroalgae have not been studied. Analysis of genomic and transcriptomic data has shown that mannitol metabolism in *S. japonica* is a circular pathway composed of four steps. In this study, one *S. japonica* mannitol-1-phosphate dehydrogenase (M1PDH2) and two mannitol-1-phosphatase (M1Pase) proteins were recombinantly expressed to analysis enzyme biochemical properties. RNA sequencing and droplet digital polymerase chain reaction were used to analyze the gene expression patterns of mannitol metabolism in different generations, tissues, sexes, and abiotic stresses. Our findings revealed insights into the mannitol synthesis pathways in brown algae. All genes were constitutively expressed in all samples, allowing maintenance of basic mannitol anabolism and dynamic maintenance of the “saccharide pool” *in vivo* as the main storage and antistress mechanism. Enzyme assays confirmed that the recombinant proteins produced mannitol, with the specific activity of SjaM1Pase1 being 1.8–4831 times that of other algal enzymes. Combined with the transcriptional analysis, SjaM1Pase1 was shown to be the dominant gene of mannitol metabolism. Mannitol metabolism genes in multicellular filamentous (gametophyte) and large parenchyma thallus (sporophyte) generations had different expression levels and responded differently under environmental stresses (hyposaline and hyperthermia) in gametophytes and sporophytes. The considerable variation in enzyme characteristics and expression of mannitol synthesis genes suggest their important ecophysiological significance in the evolution of complex systems (filamentous and thallus) and environmental adaptation of Laminariales.

## Introduction

Mannitol is one of the most widely occurring sugar alcohols in nature and is produced by a variety of living organisms, including bacteria, fungi, terrestrial plants, and algae (Iwamoto and Shiraiwa, 2005; Rousvoal *et al*., 2011). The presence of mannitol has been reported in primary endosymbiotic algae, such as those belonging to Chlorophyta (Dickson and Kirst, 1987; Dittami *et al*., 2011) and a few species of Rhodophyta (Karsten *et al*., 1997; Eggert *et al*., 2006), as well as in secondary endosymbiotic Ochrophyta algae, such as brown algae (Ji *et al*., 1980; Karsten *et al*., 1991; Gylle *et al*., 2009) and diatoms (Hellebust, 1965). As one of the main primary photosynthetic products and storage compounds in Laminariales (Kremer, 1980; Wei *et al*., 2013; Xia *et al*., 2016), mannitol can represent up to 15–26% of the dry weight of the organism (Black, 1948; Reed *et al*., 1985). Moreover, mannitol fulfills key physiological roles, including protection against environmental stress, and can act as an organic osmolyte, compatible solute, antioxidant, or thermal protectant (Iwamoto and Shiraiwa, 2005; Patel and Williamson, 2016; Tonon *et al*., 2017). Despite the importance of mannitol in the physiology of brown algae, information on its biosynthetic pathway is scarce; the functions of only a few pathway genes have been confirmed, and regulatory mechanisms are poorly understood, warranting further studies.

*Saccharina* is one of the most important macro-brown algae in the order Laminariales because of its considerably high biomass, dominance, and economic significance (Bartsch *et al*., 2008). Asian countries have been cultivating *Saccharina* species since the early 1950s (Tseng, 1987), and presently, the annual production (7.7 million tons) of this species is the second highest among all aquaculture species (FAO, 2016). Recently, compounds such as mannitol derived from *Saccharina* have been widely used in health food, medicine, analytical chemistry, and scientific research (Belcher and Nutten, 1960; Saha and Racing, 2011; Varzakas *et al*., 2012; Wakai *et al*., 2013). The life history of *Saccharina* comprises several stages, which include single-cell (meiospore), multicellular filamentous (gametophyte, n), and large parenchyma individual (sporophyte, 2n) stages (Bartsch *et al*., 2008). Its unique heteromorphic alternation of generations makes it quite different from its close relatives in the genus *Ectocarpus*, which lack the parenchyma stage (Cock *et al*., 2014). Although brown algae are the only secondary endosymbiotic taxa with sophisticated multicellularity (Knoll, 2011; Niklas and Newman, 2013; Cock *et al*., 2014), the underlying regulatory mechanisms responsible for the structural difference between filamentous brown algae (*Ectocarpus*) and heteromorphic haploid-diploid algae (*Saccharina*) are not well understood. Moreover, Laminariales, like *S. japonica*, are dominantly present in marine ecosystems of cold, temperate, and tropical coastal zones with harsh extremes (Liu and Pang, 2015). Therefore, their major photosynthetic carbohydrates, such as mannitol, may need to evolve distinctive adaptation or acclimation mechanisms.

Recently, the availability of the *Ectocarpus siliculosus* genome has paved the way for studying the molecular basis of mannitol biosynthesis in algae (Cock *et al*., 2010). The biosynthesis involves two enzymatic steps; the first step is the reduction of fructose-6-phosphate (F6P) to mannitol-1-phosphate (M1P) by mannitol-1-P dehydrogenase (M1PDH; EC 1.1.1.17), and the second step is the hydrolysis of the phosphoric ester of M1P to produce mannitol by mannitol-1-phosphatase (M1Pase; EC 3.1.3.22) (Iwamoto and Shiraiwa, 2005). Recent analysis of the distribution of mannitol biosynthesis genes in algae revealed the presence of *M1PDH* and *M1Pase* genes in these species (Tonon, 2017). There are one *M1PDH* gene, two *M1Pase* genes, and one bifunctional *M1PDH/M1Pase* fusion gene (Liberator *et al*., 1998; Groisillier *et al*., 2014); the members of Phaeophyceae, including Ectocarpales and Laminariales, possess *M1PDH* and the haloacid dehalogenases (*HAD-M1Pase*). Previous phylogenetic analyses have suggested that these genes were imported into brown algae by horizontal gene transfer from *Actinobacteria* (Michel *et al*., 2010). Later, a more comprehensive assessment across various algal lineages proved that this gene may have been present in nonphotosynthetic eukaryotic host cells involved in endosymbiosis (Tonon, 2017). Native M1PDH and M1Pase activity has previously been characterized in cell-free extracts from red algae, namely *Dixoniella grisea* (Eggert *et al*., 2006), *Caloglossa continua* (Iwamoto *et al*., 2001; Iwamoto *et al*., 2003), and *Caloglossa leprieurii* (Karsten *et al*., 1997), and from brown algae, namely *Spatoglossum pacificum, Dictyota dichotoma, Platymonas subcordiformis*, and *Laminaria digitata* (Ikawa *et al*., 1972; Grant and Rees, 1981; Richter and Kirst, 1987); however, the genes encoding these enzymes have not yet been identified. Recent structural and functional genomics research on *E. siliculosus* has analyzed mannitol metabolism, and recombinant *Ectocarpus* M1PDHcat (EsiM1PDHcat; containing only the catalytic domain) and M1Pase2 (EsiM1Pase2) have been characterized (Rousvoal *et al*., 2011; Groisillier *et al*., 2014; Bonin *et al*., 2015). Determination of kinetic parameters indicated that EsiM1PDH1cat displays higher catalytic efficiency for F6P reduction compared with M1P oxidation; EsiM1Pase2 was shown to hydrolyze the phosphate group from M1P to produce mannitol but was not active on hexose monophosphates, such as glucose-1-phosphate (G1P), glucose-6-phosphate (G6P), and F6P. Gene expression analysis showed that transcription of these three genes from *E*. *siliculosus* (filamentous brown algae) was under the influence of the diurnal cycle, and *EsM1Pase1* was highly downregulated under hyposaline stress. However, these genes are still not well understood in *S. japonica* (large parenchyma brown algae).

In this study, *M1PDH* and *M1Pase* genes and their corresponding proteins, which are involved in mannitol synthesis in *S. japonica*, were characterized. Gene expression analyses in different *Saccharina* tissue structures and samples from different stages of life cycle (including the sporophyte and gametophyte generations) and under abiotic stresses were conducted to understand the mechanisms regulating mannitol metabolism genes in Laminariales. The results of this study will expand our understanding of biosynthesis and degradation pathways and regulatory mechanism of carbohydrates and their ecophysiological and evolutionary significance in Laminariales and will provide a basis for artificial synthesis of mannitol *in vitro* and in transgenic plants for conferring salt tolerance.

## Materials and methods

### Algal sample collection

Preserved *S. japonica* haploid gametophytes (male and female gametophytes) were available as laboratory cultures and obtained from Laboratory of Genetics and Breeding of Marine Organisms. Fresh samples of the *Saccharina* sporophytes (rhizoids, stipe, blade tip, blade pleat, blade base, and blade fascia) were collected from east China (Rongcheng, Shandong Province, 37°8ʹ53ʺN, 122°34ʹ33ʺE). To detect the influences of abiotic factors, the female gametophytes and blade base of sporophytes were cultured under different temperatures (control: 8°C; hyperthermia: 18°C), salinities (control: 30‰; hyposaline: 12‰), and circadian rhythms (control: 30 μmol photons/m^2^ s for 12 h; darkness: no irradiance for 12 h). These samples were used for RNA sequencing and digital polymerase chain reaction (PCR) analysis.

### Sequence analysis

Based on the analysis of the *S. japonica* genome (NCBI accession number JXRI00000000.1, and resequencing genome data, Tao Liu, unpublished data) and transcriptome database (OneKP accession number OGZM), the unigenes related to *M1Pase* were verified using the BLASTX algorithm (http://blast.ncbi.nlm.nih.gov/Blast.cgi). Multiple sequence alignment was performed with ClustalX (Thompson *et al*., 1997). Sequence identities were calculated using the Clustal Omega tool (http://www.ebi.ac.uk/Tools/msa/clustalo/).

### Purification of recombinant proteins expressed in Escherichia coli

Genes were synthesized (Shanghai Xuguan Biotechnological Development Co. LTD) to construct recombinant plasmids. *S. japonica M1PDH1* (*SjaM1PDH1*)*, M1PDH2* (*SjaM1PDH2*), and *M1Pasel* (*SjaM1Pase1*) were cloned in pET32a and *S. japonica M1Pase2* (*SjaM1Pase2*) was cloned in pGEX-6p-1. These recombinant plasmids were transformed in *E. coli* BL21 (DE3) cells, and the integrity of their sequences was verified by sequencing. Isopropyl β-D-1-thiogalactopyranoside was added at concentrations of 0.5 mM to induce overexpression of the target proteins, and the bacterial cultures were incubated for 16 h at 20°C. His-Binding-Resin and GST-Binding-Resin were used according to the manufacturer’s instructions (www.yuekebio.com). The proteins were stored at -80°C.

### Assays for enzyme kinetics

The SjaM1PDH and SjaM1Pase activities of the purified enzymes were detected using previously described methods (Groisillier *et al*., 2014; Bonin *et al*., 2015). For enzymatic characterization, four sugar and polyol phosphoesters, which were considered potential substrates, were tested; these substrates were M1P, F6P, G1P, and G6P (Sigma, St. Louis, MO, USA). The effects of pH on the enzymatic activities of the purified proteins were determined in the range from 5.0 to 9.0 for SjaM1PDH and 5.5 to 10.5 for SjaM1Pase. The effects of temperature on these enzymes were determined over a range from 10°C to 60°C. The influence of NaCl was assessed over a final concentration range from 0 to 1000 mM in the reaction mixture. Four replicates were analyzed for each condition to ensure the consistency of the experimental results. In each case, boiled purified recombinant enzyme was used as a negative control.

### RNA sequencing and Droplet digital PCR

Total RNA was extracted using an improved CTAB method (Gareth *et al*., 2006). Three micrograms of RNA per sample was used as input material for the RNA sample preparation. Sequencing libraries were generated using NEBNext Ultra RNA Library Prep Kit for Illumina (NEB, USA) following the manufacturer’s recommendations, and index codes were added to attribute sequences to each sample. The clustering of the index-coded samples was performed on a cBot Cluster Generation System using a TruSeq PE Cluster Kit v3-cBot-HS (Illumina) according to the manufacturer’s instructions. After cluster generation, the library preparations were sequenced on an Illumina Hiseq platform, and 125-/150-bp paired-end reads were generated. HTSeq v0.6.1 was used to count the read numbers mapped to each gene. The fragments per kilobase of transcript per million mapped reads (FPKM) of each gene was then calculated based on the length of the gene and the reads count mapped to this gene. Droplet digital PCR analysis was conducted according to the previously described methods (Chi, *et al*., 2017). The results represent mean values of three replicates. All the data were subjected to one-way analysis of variance followed by Student’s *t*-tests.

## Results

### Identification of brown algal mannitol metabolism genes

Genomic sequencing data of *S. japonica* and transcriptomic data of 19 brown algal species belonging in Laminariales, Ectocarpales, Desmarestiales, Dictyotales, Fucales, and Ishigeales were identified by BLASTX analysis for mannitol metabolism genes. Two unigenes were found to be related to *M1PDHs* in most species (named *M1PDH1* and *M1PDH2* according to the naming convention for *E. siliculosus M1PDHs*), and only two Ectocarpales possessed the third *M1PDH* like *E. siliculosus* (Table S1). The identity between *M1PDH1* and *M1PDH2* within species was approximately 55.1–58.6%. *M1Pase* genes showed more conservation in brown algae. All 19 species contained two homologs of *M1Pases*, which were named *M1Pase1* and *M1Pase2*, and their identity within species was 55.6–68.2% (Table S2). Only one *M2DH* was verified in 15 Phaeophyceae species, and the identity between different species was approximately 69.9–72.8% (Table S3). All 19 species contained two homologs of HKs, which were named *HK1* and *HK2*, and their identity within species was approximately 61.0–67.4% (Table S4).

*SjaM1PDH* and *SjaM1Pase* cDNA sequences were deposited in GenBank with accession numbers MF706368, MF706369, MF440344, and MF465902. After aligning brown algal M1PDH amino acid sequences, conserved blocks A to E of PSLDRs (defined by Klimacek *et al*., 2003) were identified, while M1PDH1 and M1PDH3 had an additional extension N-terminal domain (Figure 1A). The comparison of M1Pases confirmed the conservation of brown algal M1Pases, including the catalytic machinery and the Mg^2+^ cofactor binding site (Figure 1B).

**Figure 1.**
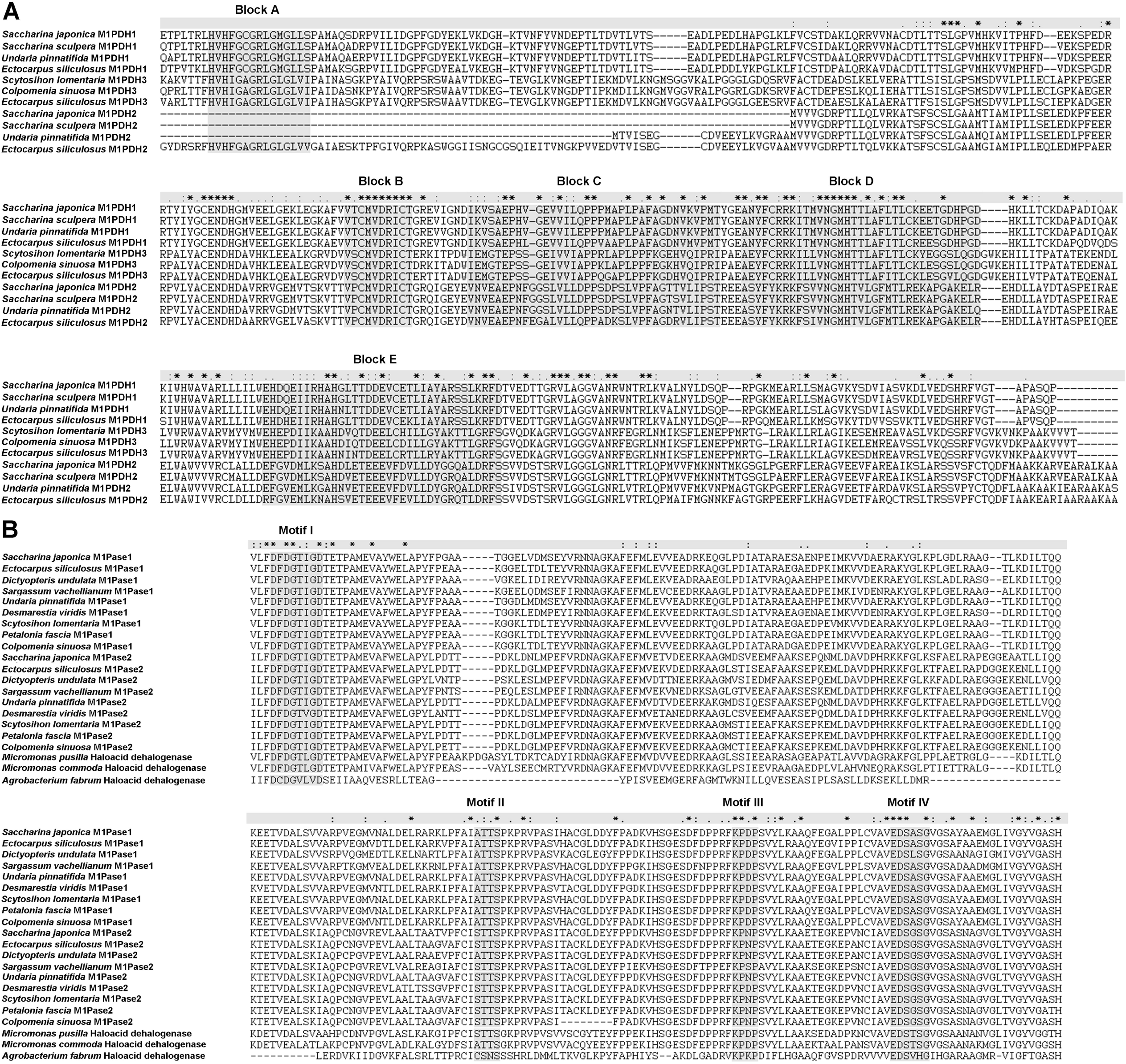
Structure-based sequence alignment of SjaM1PDHs (A) and SjaM1Pases (B). A. Alignment of the M1PDH from *Saccharina japonica* and some other brown algae species. Five conserved blocks as defined by Klimacek *et al*. (2003) for PSLDRs, named A to E, are represented above the conserved consensus sequence. B. Alignment of the crystallized HAD-like protein ATU0790 from *Agrobacterium tumefaciens* strain C58 (PDB code 2FDR) with orthologs of M1Pases from different organisms. Motifs I to IV, defined according to Burroughs *et al*. (2006), are represented above the conserved consensus sequence.

### Characterization and confirmation of the functions of mannitol biosynthesis genes from S. japonica

Native *SjaM1PDH2, SjaM1Pase1*, and *SjaM1Pase2* were overexpressed in *E. coli* to characterize the M1PDH and M1Pase activity of *S. japonica*. Although several attempts were made to overexpress *SjaM1PDH1*, no protein was observed, similar to *E. siliculosus M1PDH1*. The specificity of SjaM1PDH2 was determined by assaying activity in presence of different potential substrates. Reduction of F6P, G6P, and G1P and oxidation of M1P, F6P, G6P, and G1P were tested. These experiments showed that the SjaM1PDH2 enzyme only had reduction activity in the direction of mannitol synthesis; no oxidation activity was detected (Table 1). In addition, the reduction activity was also detected for other sugar substrates, indicating that SjaM1PDH2 was not specific for F6P (Table 2). Purified SjaM1PDH2 had a specific activity of 0.36 μmol/mg protein/min for F6P reduction with NADH at pH 8.0. This activity was in the range of those measured for algal M1PDHs listed in Table 1. The SjaM1Pase activities were determined in 100 mM Tris-HCl buffer. The specific activity of SjaM1Pase1 (144.93 μmol/mg protein/min) was significantly higher (almost 22 times) than that of SjaM1Pase2 (6.60 μmol/mg protein/min) in the presence of 1 mM M1P (Table 3). The enzymatic reactions were performed in the presence of F6P, G1P, and G6P to investigate the substrate specificity of SjaM1Pases. The phosphatase activity was also detected for all sugar phosphates that were tested for each *Saccharina* protein. The activity of SjaM1Pase1 for such substrates was always lower than for M1P, as was observed for most brown and red algae M1Pases (Table 4). However, SjaM1Pase2 exhibited the highest phosphatase activity in the presence of G1P, which was almost 1.1 times that of M1P, and was similar to the substrate specificity of M1Pase from *Dixoniella grisea*. In addition, more than 90% enzymatic activity was detected in SjaM1PDH2 and both SjaM1Pases after storage at 4°C for 72 h, suggesting that the recombinant proteins were quite stable under the purification conditions tested.

**Table 1.** Comparison of the biochemical characterization of M1PDH and its activity determined in brown and red algae.

| | Specific activity<br>/$V_{max}$ ($\mu\text{mol mg}$<br>protein<sup>-1</sup> min<sup>-1</sup>) | $K_m$<br>(mM)<br>M1P | $k_{cat}$<br>(s<sup>-1</sup>) | pH<br>optimum | Temp<br>optimum<br>(°C) | References |
| --- | --- | --- | --- | --- | --- | --- |
| **Brown algae** |  |  |  |  |  |  |
| *Saccharina japonica*<br>(SjaM1PDH2) | 0.36 | 2.02 | 0.44 | 8.0 | 40 | This study |
| *Ectocarpus siliculosus*<br>(EsM1PDH1cat) | 46.2 | 0.19 | 32.3 | 7.0 | 30 | Bonin *et al.*, 2015 |
| *Spatoglossum pacificum* | 0.46 | 0.28 | NT | 6.5 | NT | Ikawa *et al.*, 1972 |
| *Platymonas subcordiformis* | NT | 5.5 | NT | 7.5 | NT | Richter and Kirst,<br>1987 |
| **Red algae** |  |  |  |  |  |  |
| *Caloglossa continua* | 228 | 0.15 | NT | 7.2 | NT | Iwamoto *et al.*, 2003 |
| *Caloglossa leprieurii* | 0.26 | 1.4 | NT | 7.0 | NT | Karsten *et al.*, 1997 |
| *Dixoniella grisea* | 0.06 | 1.18 | NT | NT | NT | Eggert *et al.*, 2006 |

NT: Not tested.

**Table 2.**
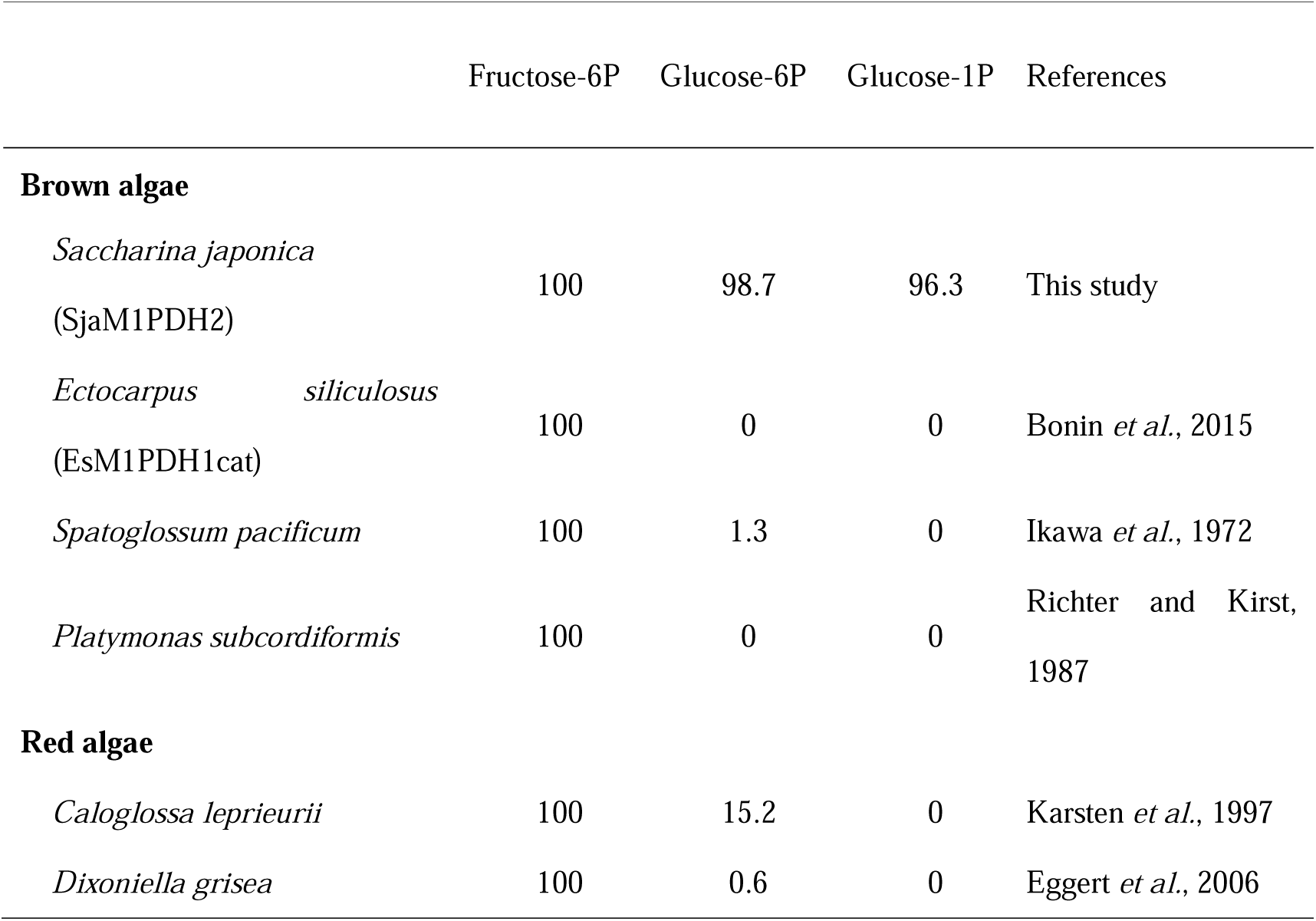
Summary of M1PDH activity determined in brown and red algae. Percentages of maximum activities were calculated according to values reported in the different publications indicated below.

|  | Fructose-6P | Glucose-6P | Glucose-1P | References |
| --- | --- | --- | --- | --- |
| **Brown algae** |  |  |  |  |
| *Saccharina japonica*<br>(SjaM1PDH2) | 100 | 98.7 | 96.3 | This study |
| *Ectocarpus siliculosus*<br>(EsM1PDH1cat) | 100 | 0 | 0 | Bonin *et al.*, 2015 |
| *Spatoglossum pacificum* | 100 | 1.3 | 0 | Ikawa *et al.*, 1972 |
| *Platymonas subcordiformis* | 100 | 0 | 0 | Richter and Kirst,<br>1987 |
| Red algae |  |  |  |  |
| *Caloglossa leprieurii* | 100 | 15.2 | 0 | Karsten *et al.*, 1997 |
| *Dixoniella grisea* | 100 | 0.6 | 0 | Eggert *et al.*, 2006 |

**Table 3.**
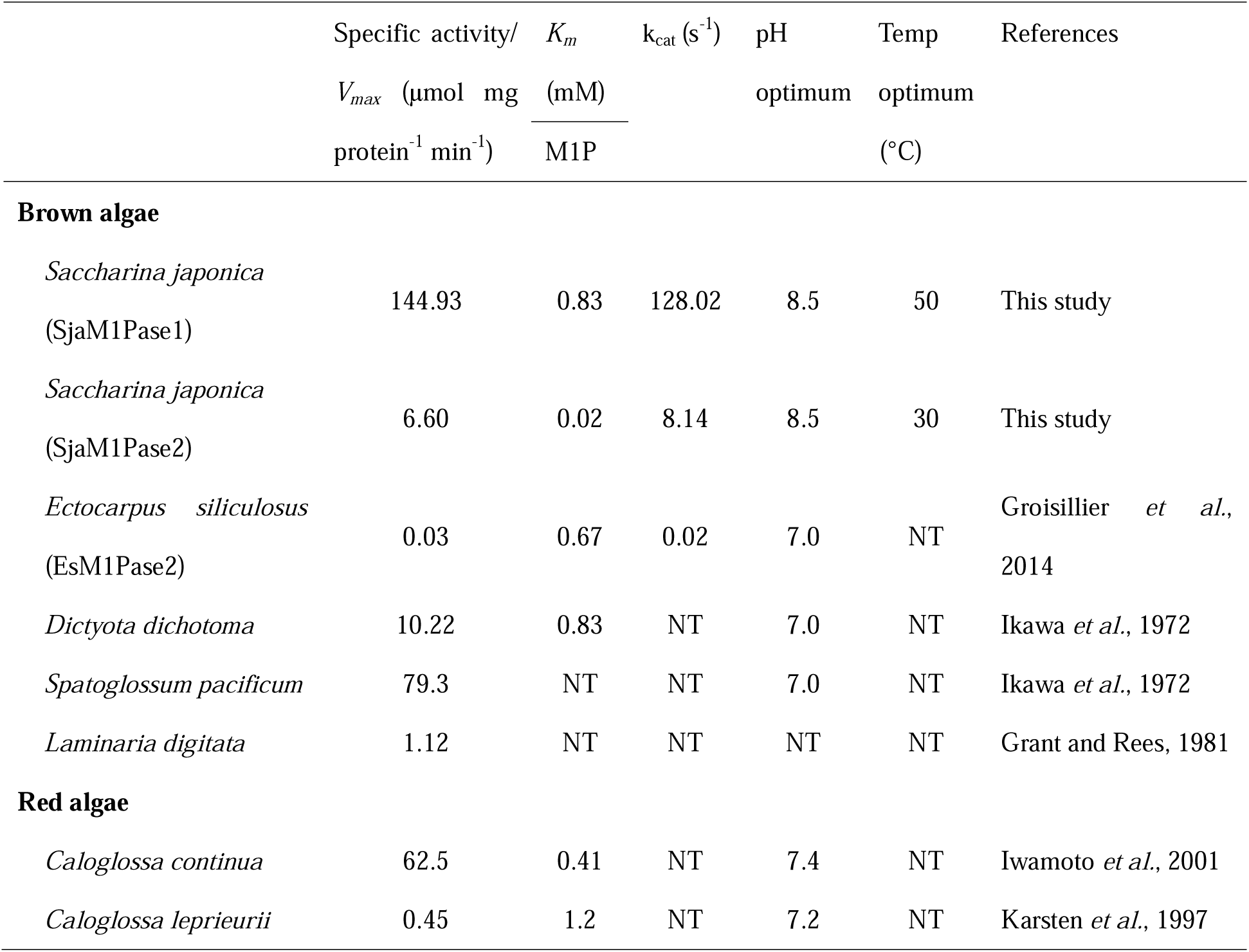
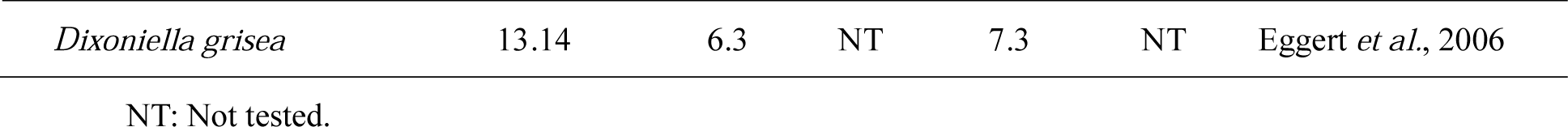
Comparison of the biochemical characterization of M1Pase and its activity determined in brown and red algae.

**Table 4.**
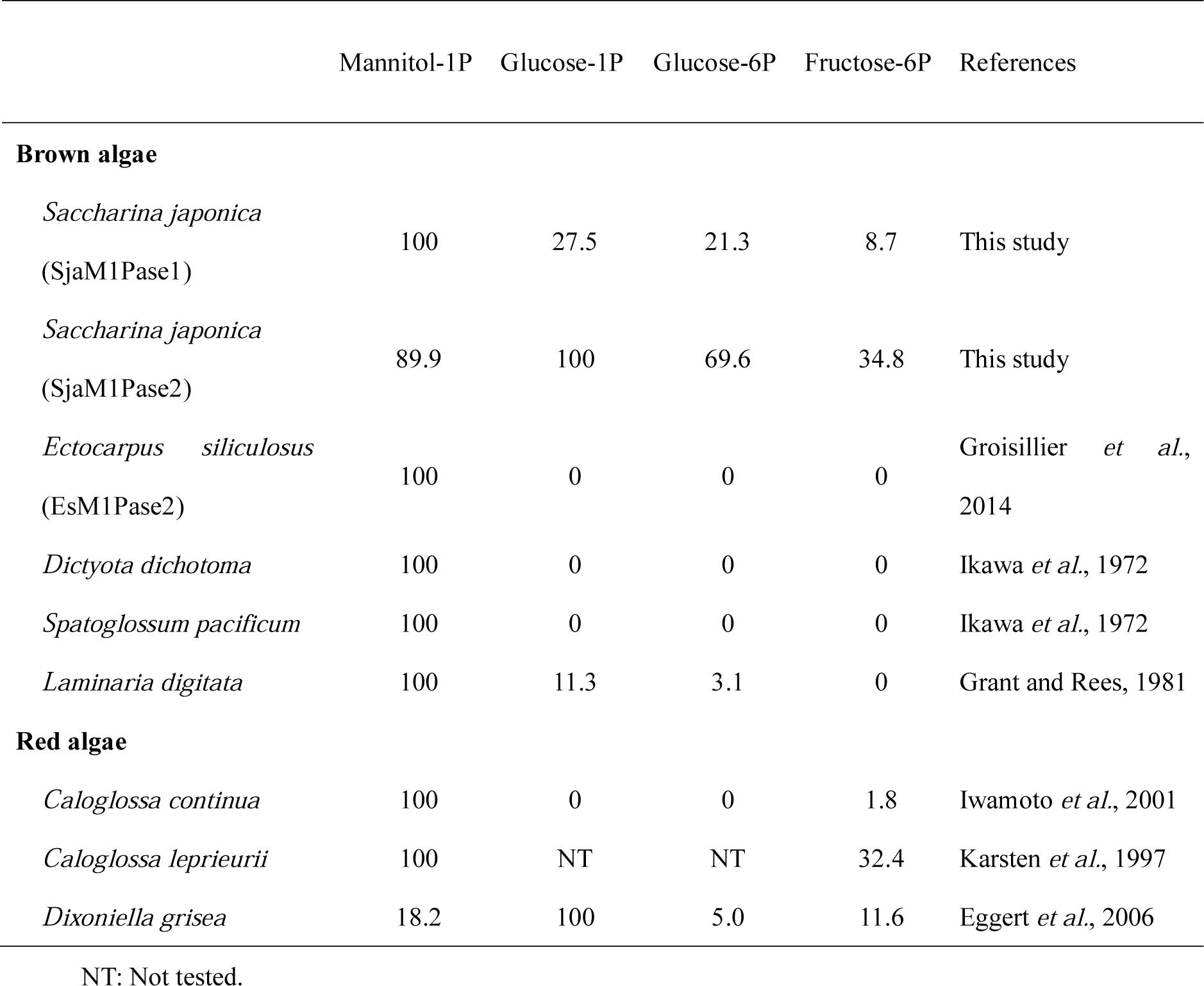
Summary of M1Pase activity determined in brown and red algae. Percentages of maximum activities were calculated according to values reported in the different publications indicated below.

The activities of the three purified proteins under different conditions of temperature and pH were determined to elucidate the biological characteristics of mannitol synthesis enzymes. Based on these experiments, the optimum temperature for SjaM1PDH2 was found to be 40°C, whereas the activities were 82% and 91% of the maximum activity at 30°C and 50°C. The optimum temperature for SjaM1Pase1 was 50°C, with 95% and 83% of residual activity at 40°C and 60°C, respectively. The optimum temperature for SjaM1Pase2 was much lower (30°C), with the activity being less than 54% of the maximum activity at other temperatures (Figure 2A–C). The optimum pH for SjaM1PDH2 was determined to be 8.0, with 51% and 56% of residual activity at pH 7.0 and 9.0, respectively. The optimum pH for both SjaM1Pase1 and SjaM1Pase2 was determined to be 8.5, and 78% of the SjaM1Pase2 activity remained intact at pH values from 7.5 to 9.5, whereas the corresponding activities for SjaM1Pase1 were less than 72% of the maximum activity (Figure 2D–F).

**Figure 2.**
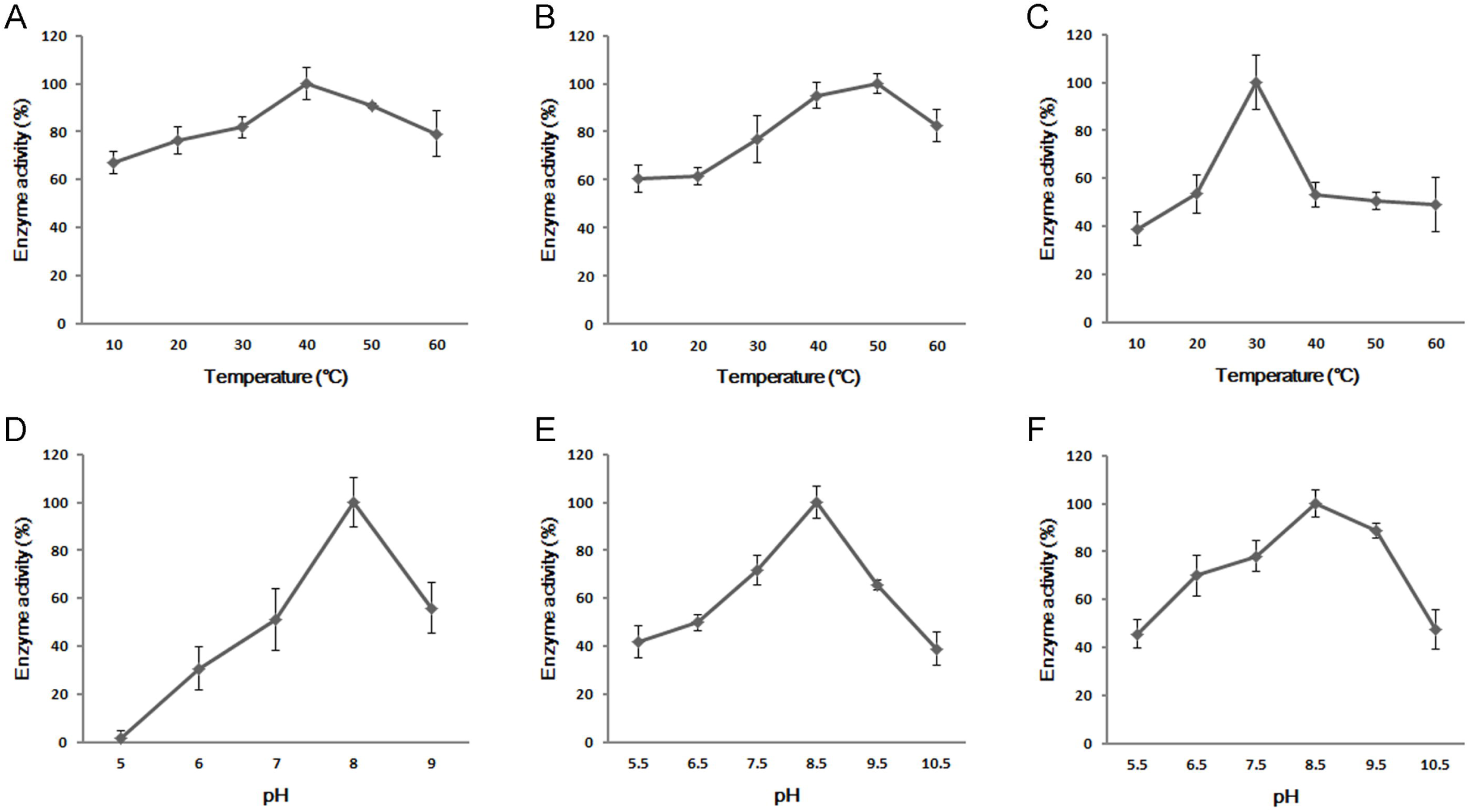
Enzymatic characteristics of recombinant SjaM1PDH2 and SjaM1Pases. A. Influence of temperature on SjaM1PDH2 activity. Enzyme activity at 40°C was set to 100%. *P* < 0.05 compared to 40°C. B. Influence of temperature on SjaM1Pase1 activity. Enzyme activity at 50°C was set to 100%. *P* < 0.05 compared to 50°C. C. Influence of temperature on SjaM1Pase2 activity. Enzyme activity at 30°C was set to 100%. *P* < 0.01 compared to 30°C. D. Influence of different pH on the activity of SjaM1PDH2. Enzyme activity at pH 8.0 was set to 100%. *P* < 0.01 compared to pH 8.0. E. Influence of different pH on the activity of SjaM1Pase1. Enzyme activity at pH 8.5 was set to 100%. *P* < 0.01 compared to pH 8.5. F. Influence of different pH on the activity of SjaM1Pase2. Enzyme activity at pH 8.5 was set to 100%. *P* < 0.05 compared to pH 8.5. The values represent means ± SD calculated from three assays.

The purified recombinant SjaM1PDH2 and SjaM1Pase proteins exhibited typical Michaelis-Menten kinetics when assayed with increasing concentrations of their substrates, and apparent *V_max_* and *K_m_* values were determined from the Lineweaver-Burk plots (Figure 3). The *K_m_* value for SjaM1PDH2 was 2.02 mM, which was about 10-fold higher than that for EsiM1PDH1cat (0.19 mM), indicating lower substrate binding capacity than EsiM1PDH1cat (Table 1). The substrate binding capacity of two *S. japonica* M1Pases showed large differences: the binding capacity of SjaM1Pase2 (*K_m_* = 0.02 mM) was highest (almost 21–315 times) among the M1Pases of other brown and red algae, almost 41 times higher than that of SjaM1Pase1 (*K_m_*= 0.83 mM). Interestingly, both the SjaM1Pases showed much higher catalytic reaction efficiency and catalytic rates than EsM1Pase2 (Table 3); SjaM1Pase1 had the highest k_cat_ value (almost four-orders of magnitude higher than *Ectocarpus* M1Pase2).

**Figure 3.**
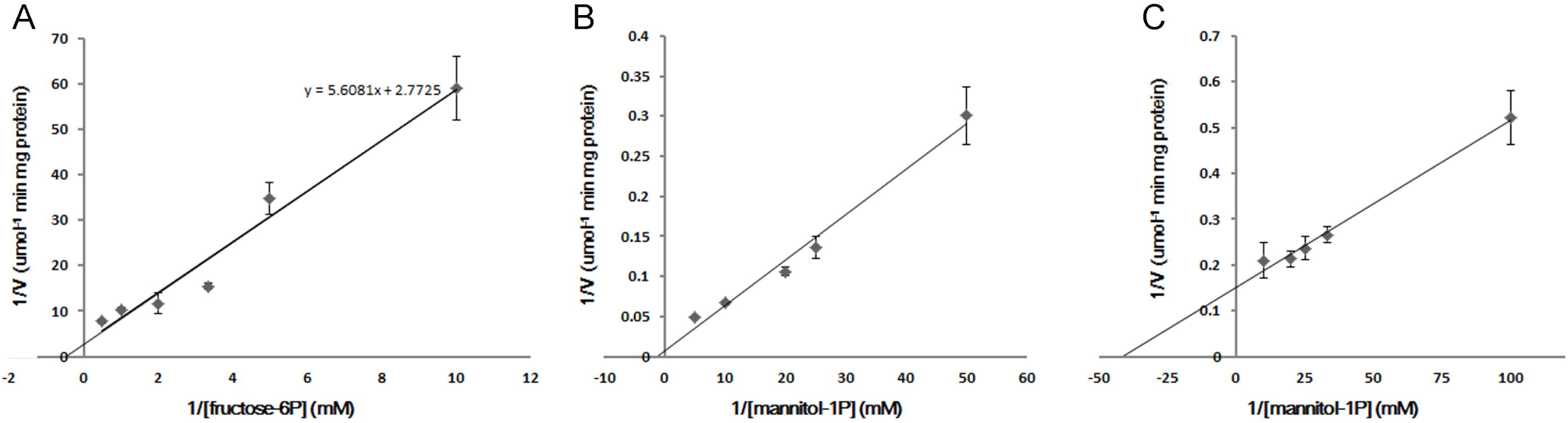
Kinetics of enzymatic activity of SjaM1PDH2 (A), SjaM1Pase1 (B) and SjaM1Pase2 (C). The values represent means ± SD calculated from three assays.

The assays conducted to assess the influence of NaCl concentration on the activity of the recombinant *S. japonica* proteins showed a dose-dependent effect on SjaM1Pases (Figure 4A, B). A nearly linear decrease in activity was observed in the presence of NaCl at concentrations ranging from 0 to 1 M. About 60% of the initial activity remained intact in the presence of 1 M NaCl for both the enzymes, which indicated that these SjaM1Pases may resist high NaCl concentrations. However, SjaM1PDH2 activity was not affected by NaCl addition (Figure 4C).

**Figure 4.**
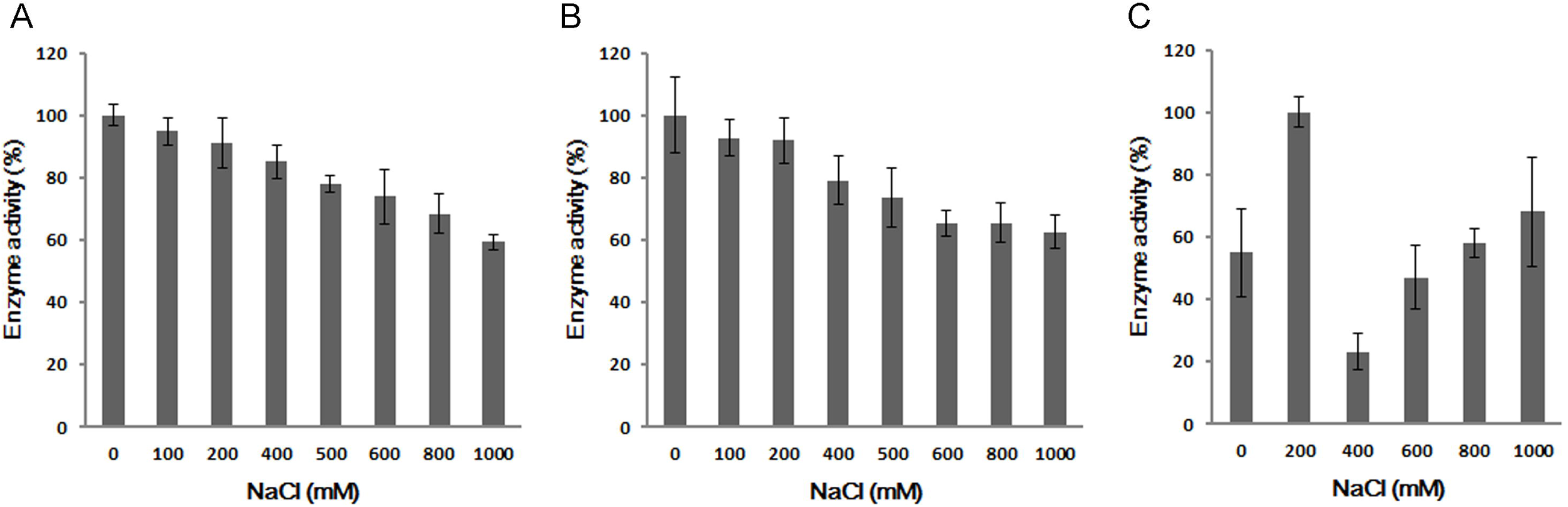
Influence of NaCl concentration on the activities of recombinant SjaM1Pase1 (A), SjaM1Pase2 (B) and SjaM1PDH2 (C). The values represent means ± SD calculated from three assays.

### Transcriptiomic analysis in mannitol metabolism

The expression of mannitol synthesis genes from *S. japonica* was determined by analyzing the transcriptomic data of *S. japonica* (Tao Liu, unpublished data), as well as the transcriptomic data of 19 Phaeophytes published by us in the OneKP database (www.onekp.com). All the Phaeophyceae species expressed two *SjaM1Pase* genes, and more than 80% of Phaeophyceae species expressed two *SjaM1PDH* genes.

To analyze the gene expression levels in different generations, tissues, sexes, and environmental conditions of *S. japonica*, RNA-seq and droplet digital PCR experiments were conducted. The regulation of metabolism in *Saccharina* is highly complex and can be divided into four general levels. First, the genes (including different gene family numbers) were all expressed constitutively. The gene expression levels were examined at various generations (female and male gametophytes; sporophytes) and in different tissues (rhizoids, stipe, blade tip, blade pleat, blade base, and blade fascia) of *Saccharina*. All four genes (including seven family numbers) were detected in different samples (Figure 5). The FPKM value range of *Saccharina* transcripts was approximately 1.2–300 (Table 5).

**Figure 5.**
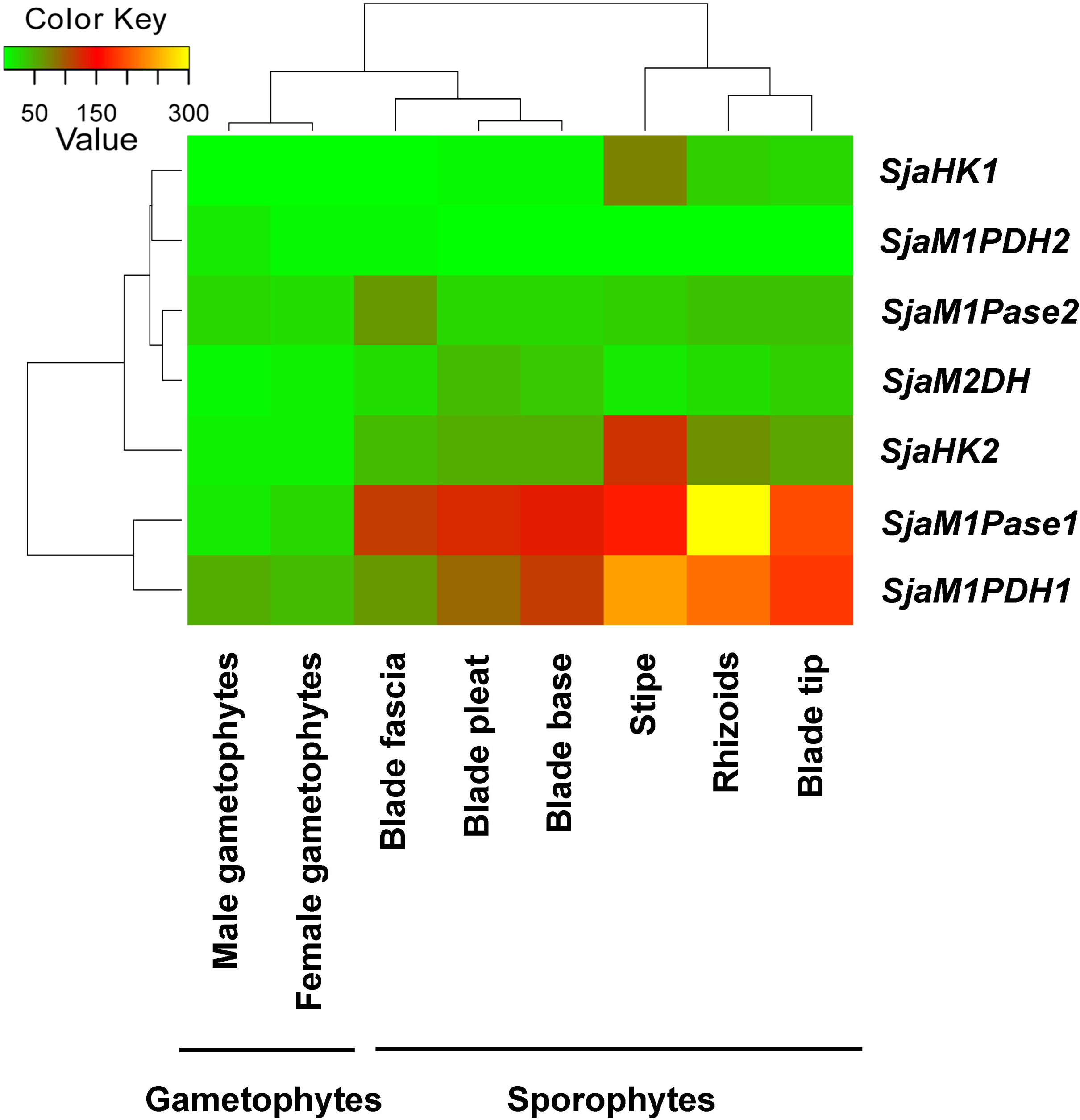
Expression levels of mannitol metabolism genes in different generations (sporophytes and gametophytes) and tissues (rhizoids, stipe, blade tip, blade pleat, blade base, and blade fascia). All genes were constitutive expressed in different samples.

**Table 5.** The upregulated and downregulated ratios of the gene FPKM doses under hyposaline (12‰) and hyperthermia (18°C) conditions.

| Genes | Sporophytes |  | Gametophytes |  |
| --- | --- | --- | --- | --- |
|  | Control/<br>Hyposaline | Control/<br>Hyperthermia | Hyposaline/<br>Control | Hyperthermia/<br>Control |
| *SjaM1PDH1* | 2.9 | 2.0 | 3.7 | 1.9 |
| *SjaM1PDH2* | 3.7 | 0.2 | 1.2 | 7.9 |
| *SjaM1Pase1* | 1.9 | 3.0 | 1.9 | 2.7 |
| *SjaM1Pase2* | 3.5 | 0.5 | 2.0 | 1.2 |
| *SjaM2DH* | 2.7 | 1.7 | 16.2 | 12.5 |
| *SjaHK1* | 3.6 | 2.9 | 4.6 | 1.3 |
| *SjaHK2* | 3.5 | 1.4 | 6.6 | 2.3 |

Second, most pathway genes encode reversible enzymes, which control the balance between mannitol and F6P and dynamically maintain the “saccharide pool” *in vivo*. As a circular pathway, mannitol metabolism is closely related to alginate, fucoidan, laminarin, and trehalose metabolism through the intermediate product F6P. The first gene that transforms F6P to mannitol is *M1PDH* (*M1PDH1* and *M1PDH2*), and the first gene that transforms into alginate and fucoidan is mannose phosphate isomerase *MPI* (*MPI1* and *MPI2*) (Chi *et al*., 2017). The expression levels of *SjaM1PDH* and *SjaMPI* were opposite in most tissues (Figure 6). For example, the expression level of *SjaMPI1* was 3.8 times that of *SjaM1PDH1* in blade fascia, whereas the expression level of *SjaM1PDH1* was 2.7 times that of the *SjaMPI1* in stipe.

**Figure 6.**
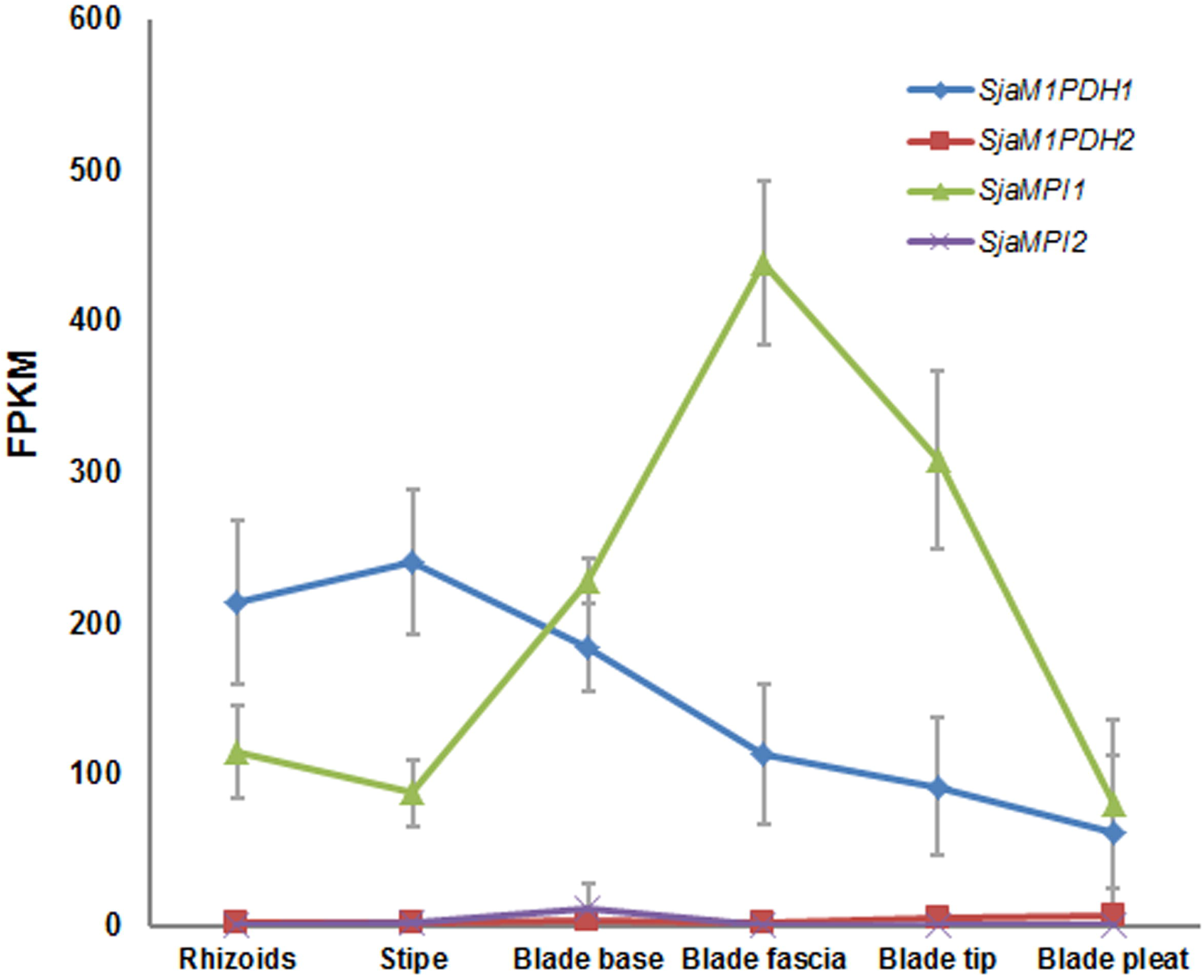
Expression levels of *SjaM1PDHs* and *SjaMPIs* in different tissues (rhizoids, stipe, blade tip, blade pleat, blade base, and blade fascia). The expression levels of *SjaM1PDHs* and *SjaMPIs* were opposite in most tissues.

Third, enzyme activity and gene expression analyses were used to determine the dominant genes in mannitol synthesis. The mannitol synthesis pathway includes two genes, *M1PDH* and *M1Pase*; the former catalyzes the reversible reaction, and the latter only catalyzes the positive reaction. Regarding the two *M1Pase* family members, the expression of *SjaM1Pase1* in different tissues was significantly higher than that of *SjaM1Pase2* (the former was 2.0–19.4 times that of the latter; Figure 7). The expressional levels of *SjaM1Pase2* in different tissues were not significantly different, but were adversely affected by *SjaM1Pase1* under hyperthermia (18°C) stress. *SjaM1Pase2* was obviously upregulated 2.0 times, whereas *SjaM1Pase1* was downregulated 2.0 times.

**Figure 7.**
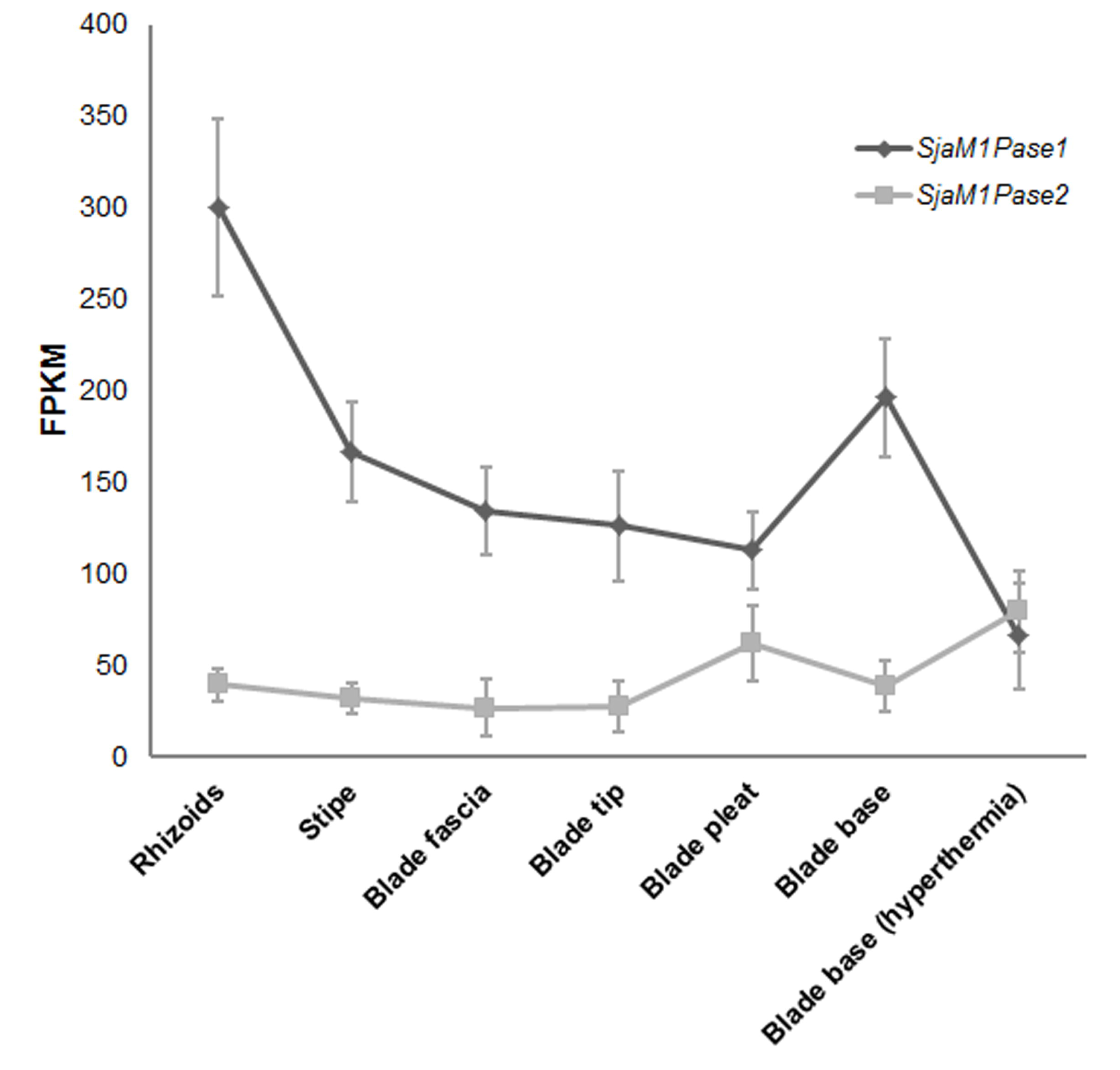
Expression levels of *SjaM1Pases* in different tissues (rhizoids, stipe, blade tip, blade pleat, blade base, and blade fascia) and under hyperthermic stress. The expression of *SjaM1Pase1* was higher than that of *SjaM1Pase2* in all tissues. The expressional level of *SjaM1Pase2* was adversely affected by *SjaM1Pase1* under hyperthermia (18°C) stress.

Finally, the overall gene expression profiles differed between the gametophyte and sporophyte generations. The average gene expression levels in the gametophyte and sporophyte generations were further compared. Most genes (*SjaM1PDH1*, *SjaM1Pase1, SjaM2DH, SjaHK1*, and *SjaHK2*) were expressed at significantly higher levels in sporophytes than in gametophytes, with increases of 2.8-, 7.3-, 3.1-, 5.6-, and 4.7-fold, respectively (Figure 8A). However, the expression levels of *SjaM1PDH2* and *SjaM1Pase2* did not differ significantly between the two generations. In addition, there were no significant differences in the expression of all pathway genes in female and male gametophytes (Figure 8B). To complete these observations, further analysis was focused on changes in the expression of the genes using algal samples subjected to abiotic stress. Interestingly, these genes in different generations had the opposite response mechanisms to environmental stress. Under hyposaline conditions, the transcript levels of all genes were upregulated, exhibiting increases of 1.2–16.2-fold in gametophytes. In contrast, the expression levels of all genes were decreased 1.9–3.7-fold in sporophytes (Table 5). Furthermore, these variations followed a similar trend under hyperthermic stress. The transcript levels of all genes were upregulated (1.2–12.5 times) in gametophytes, whereas most genes (except *SjaM1PDH2* and *SjaM1Pase2*) were downregulated (1.4–3.0 times) in sporophytes (Table 5).

**Figure 8.**
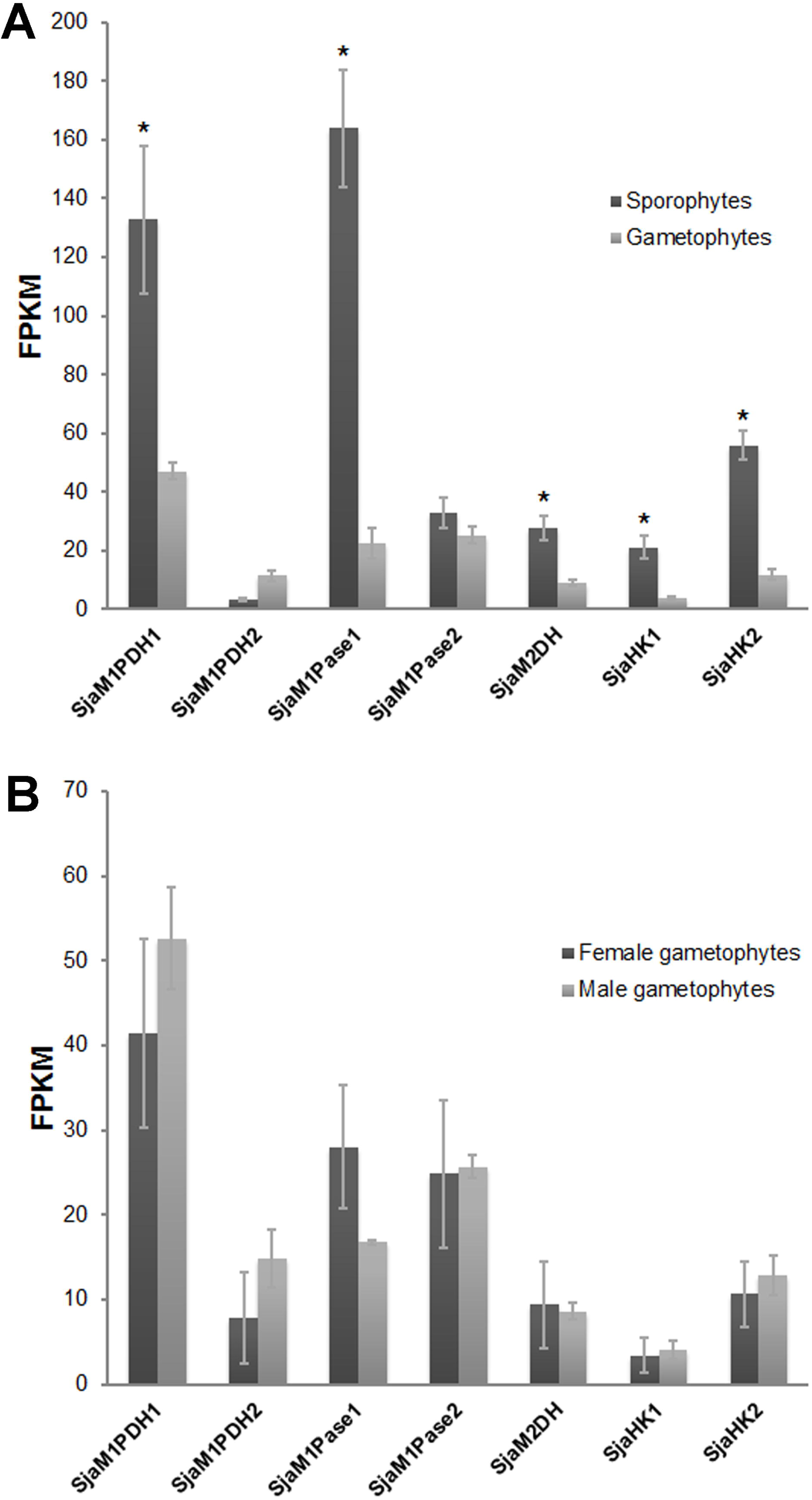
(A) Expression levels of mannitol metabolism genes at different generations (gametophytes and sporophytes). Most of the gene transcript levels of sporophytes were much higher than that of gametophytes (*P* < 0.05). (B) Expression levels of mannitol metabolism genes at different sexse (female gametophytes and male gametophytes). No significant differences were observed between these samples.

## Discussion

### Mannitol metabolism genes were constitutively expressed in brown algae, which satisfied the requirement for mannitol synthesis and accumulation for physiological metabolism

Mannitol is the fundamental carbon-storage molecule and osmotic regulator in brown algae, and mannitol metabolism is one of the main traits that makes brown algae unique compared with other eukaryotic algae (Shao *et al*., 2014). As a basal metabolic pathway, mannitol metabolism has fewer pathway steps (a total of four steps) and relatively few gene family members (1–2 copies of each gene in most brown algae). This mechanism differs obviously from some special elements, such as halogen metabolism (about 89 gene family members in *S. japonica*, unpublished data), and because of the single product (mannitol), this pathway does not contain complex synthesis and modification genes, such as glycosyltransferase, sulfurtransferase, and mannuronate C5-epimerases genes, as observed in alginate and fucoidan metabolism (which contain dozens to more than 100 genes) (Chi *et al*., 2017). Therefore, analysis of mannitol metabolism is critical for the study of pathway regulation and environmental adaptation of brown algae.

Of the four pathway genes, *M1PDH, M1Pase*, and *HK* genes have a “backup” gene (only *M2DH* has one copy). Moreover, about 80% of brown algal species in our analysis also harbor similar gene numbers. There are three *M1PDH* unigenes in *E. siliculosus*, but only two copies are found in the *S. japonica* genome. The transcriptomic data of 19 Phaeophyceae species also identified two *M1PDH* copies expressed in brown algae, except Ectocarpales. To confirm the *M1PDH* gene duplicates in brown algae, genome sequencing data of *Undaria pinnatifida* and *Costaria costata* (both belonging to Laminariales) were analyzed; these organisms were both found to possess *M1PDH1* and *M1PDH2*. The third unigene of *M1PDH* in Ectocarpales may be explained by the duplication of an *M1PDH1* sequence (Tonon *et al*., 2017). Unlike in other primary endosymbiotic (e.g., red algae) and secondary endosymbiotic (e.g., Dictyochophyceae) algae, which have only one gene copy (Tonon *et al*., 2017), brown algae have two *M1Pase* genes (Table S2). Transcriptome analysis demonstrated that all four genes were constitutively expressed (at least at the RNA level) in brown algae. In addition, further investigations revealed that these genes are also constitutively expressed at various generations and in different tissues (Figure 5; Table S5). Even under dark conditions, all genes in the pathway were detected (the FPKM dose was 12.0–158.8, Table S5). During the development of *Saccharina*, zygotes divide continuously from a single cell to form thallus sporophytes, which exhibit consistent increases in length, width, and thickness. Mannitol is a central compound in plant carbon metabolism and in transportation and distribution of the organic assimilate. Moreover, mannitol has important physiological functions, such as osmotic regulation, antioxidant, thermal protection, and respiration substrate (Schmitz, *et al*., 1972; Davison & Reed, 1985). Therefore, these results suggest that brown algae consistently synthesize mannitol for carbon storage and energy.

### Specific expression and regulation of the circular metabolism

*Saccharina* mannitol metabolism is a unique circular metabolic pathway, having few steps (four genes, seven copies), i.e., far less than other metabolic pathways such as alginate (six genes, 133 copies) (Chi *et al*., 2017), fucoidin (nine genes, 71 copies) (Chi *et al*., 2017), and trehalose (three genes, 11 copies) (Chi *et al*., 2015; unpublished data) pathways. As the shared substrate, F6P can be used to synthesize other basic metabolites, such as alginate and fucoidan, and the regulation of mannitol synthesis genes in Laminariales could involve a complex integrated system. For example, the expression level of the first gene in mannitol, alginate, and fucoidan metabolic pathways was found to differ substantially among tissues (Figure 6), indicating that more F6P could be utilized in the synthesis of alginate and fucoidan. This is consistent with the finding that the accumulation of mannitol has an inverse relationship with that of alginate and fucoidan (Kaliaperumal and Kalimuthu, 1976; Ji, 1963). Mannitol metabolism can finely regulate the balance between mannitol and F6P and further affect other related pathways, such as alginate and fucoidan, which function to maintain the “saccharide pool” *in vivo*.

### The analysis of enzyme activity and gene expression reveals that M1Pase1 is the dominant gene of mannitol synthesis

As the enzyme catalyzing the positive reaction, M1Pase may be the rate-limiting gene for mannitol synthesis. Although the two copies of M1Pase were found to be expressed constitutively, there were major differences in their enzyme activities and expression patterns.

M1Pase enzymes exhibit mannitol biosynthesis activity and have different biochemical properties. In brown algae, only one homolog of the *M1Pase* gene (*M1Pase2*) was confirmed to show activity in *E. siliculosus* (Groisillier *et al*., 2014; Bonin *et al*., 2015). No enzymatic studies have been conducted for *M1Pase1*; only their nucleotide sequences were reported from *E. siliculosus*. In this study, SjaM1Pase1 and SjaM1Pase2 were both confirmed to have M1Pase activity and were assumed to be involved in the mannitol biosynthesis pathway in brown algae (Figure 2). The specific enzyme activity of SjaM1Pase1 was much higher (21.9 times) than that of SjaM1Pase2, and the former enzyme had a higher k_cat_ value (15.7 times) than that of the latter (Table 3). This indicated that SjaM1Pase1 had stronger catalytic ability than SjaM1Pase2. In terms of expression levels, SjaM1Pase1 also showed significantly higher expression than SjaM1Pase2 in all tissues (Figure 7). These results suggested that *SjaM1Pase1* was the dominant gene in mannitol synthesis and was mainly responsible for the synthesis and accumulation of mannitol, whereas *SjaM1Pase2* acted as a backup, having an important role under some conditions. For example, when *SjaM1Pase1* was downregulated under hyperthermic stress in sporophytes, *SjaM1Pase2* exhibited the opposite response mechanism under the same temperature stress.

All genes and copies (family members) of mannitol metabolism in *S. japonica* are involved in mannitol metabolism and show the characteristic of higher expression for the dominant gene. Multiple copies of genes are important for maintaining the normal biochemical metabolism in evolution. Mannitol metabolism gene duplication is an important biological mechanism to maintain the core photosynthetic carbon storage in *S. japonica*. Different copies of genes were still transcribed, and their gene products exhibited activity conducive to avoiding the influence of gene loss, recombination, and mutation on gene integrity and enzyme function. Interestingly, most genes were downregulated under hyperthermic stress in sporophytes, whereas *SjaM1PDH2* and *SjaM1Pase2* were upregulated (about 5.6 and 2.0 times, respectively; Figure 7, Table S5). These findings suggested that there were different mechanisms of regulation for backup genes.

### Complex regulation of the expression of mannitol metabolism-related genes revealed the importance of this process in the evolution (filamentous and thallus) and environmental adaptation of brown algae

Laminariales algae have a heteromorphic haploid-diploid life cycle, with a macroscopic thallus sporophyte and microscopic gametophyte among different generations (Bartsch *et al*., 2008). Most mannitol metabolism genes (except *SjaM1PDH2* and *SjaM1Pase2*) had significantly higher expression in sporophytes (filamentous generation) than in gametophytes (thallus generation; Figure 8A). This is consistent with the results of our mannitol content analysis in *Saccharina*, in which the dry weight of mannitol was found to be lower in gametophytes (male gametophytes with 23.4% and female gametophytes with 24.6%) than in sporophytes (26.3%). These results indicated that the sporophytes may have much higher ability to synthesize mannitol, potentially because the sporophytes (large thallus with tissue differentiation) require more mannitol synthesis and degradation to satisfy normal growth and development and adapt to the changing environment. However, there were no significant differences in gene expression between female and male gametophytes (Figure 8B). This is also consistent with our observation that the mannitol contents in male and female gametophytes were 23.4% and 24.6%, respectively, suggesting that different sexes of gametophytes had similar mechanisms for regulating mannitol metabolism.

The mechanisms regulating mannitol metabolism in response to environmental stress are opposite between different generations. Under hyposaline stress, all genes in gametophytes were upregulated, consistent with the results observed in *M1Pase2* from *E. silicullosus* (Groisillier *et al*., the 2014), whereas the sporophyte genes were all downregulated (Figure 9A). Similar results were observed under hyperthermic conditions (Figure 9B). Notably, mannitol is the main organic osmolyte in *Saccharina* and most other brown algae, countering salinity stress and acting as an antioxidant and heat protectant (stabilization of proteins) (Schmitz *et al*., 1972; Davison and Reed, 1985; Iwamoto and Shiraiwa, 2005). The gametophyte stage is the stage that is most vulnerable to external stress in the entire life-history of the organism (Ye *et al*., 2015). Therefore, during this stage, mannitol metabolism is increased to respond to stress, particularly as an osmotic adjustment substance to quickly respond to hyposaline stress, maintain intracellular osmotic pressure, and protect algae against hyperthermia-induced damage. Distinct regions of carbon sources and carbon sinks exist along the thalli because of their large size and differentiation (Schmitz and Lobban, 1976; Buggeln, 1983). *Saccharina* involves transport from source to sink, i.e., from mature blade areas, which produce a surplus of photoassimilates, to the intercalary carbon-requiring meristems (Bartsch *et al*., 2008). The imported organic compounds in the sink tissues are rapidly metabolized and incorporated into polysaccharides and proteins (Schmitz and Lobban, 1976). The blade base of the sporophytes (meristem) exhibited reduced mannitol metabolism, probably related to reduction of mannitol degradation and incorporation, thereby decreasing transportation from carbon sources to carbon sinks and maintaining mannitol accumulation in mature blades to adapt to stress.

### Comparative analysis of RNA-seq and droplet digital PCR

The droplet digital PCR method was used to verify the transcriptional sequencing results. The gene expression in gametophytes was analyzed under stresses (hyposaline and hyperthermia). Most of the genes were upregulated under these stress conditions, consistent with the results of RNA-seq. For example, *SjaM1PDH1* increased by 4.0 and 1.3 times under hyposaline and hyperthermia stresses, respectively, as determined by droplet digital PCR analysis, whereas RNA-seq analysis showed that *SjaM1PDH1* increased by 3.7 and 1.9 times under the same conditions (Table S6). RNA-seq and droplet digital PCR analyses yielded similar experimental results. Notably, droplet digital PCR was relatively inexpensive and had a shorter experimental time, whereas RNA-seq was beneficial for further analysis of the relationship between mannitol metabolism and other pathways.

### Characterization of mannitol synthesis genes in brown algae may provide more effective genes for industrial production of mannitol and for plant genetic breeding

Mannitol is a commercially valuable compound and is now widely used in the food, pharmaceutical, medical, and chemical industries (Saha and Racine, 2011; Song and Vieille, 2009). Most of the commercial production of mannitol is carried out by chemical hydrogenation of fructose, or it is extracted from seaweed (Saha and Racine, 2011; Xia *et al*., 2016). Because of the problems associated with chemical production and extraction, microbial production has been the subject of significant interest in recent years (De Guzman, 2005; Saha and Racine, 2011). The most widely used *M1Pase* gene from a protozoan parasite *Eimeria tenella* (Liberator *et al*., 1998) had been expressed in cyanobacterium (Jacobsen and Frigaard, 2014), proteobacteria (Reshamwala *et al*., 2014), and firmicutes (Wisselink *et al*., 2005) and resulted in the accumulation of mannitol in the cells and in the culture medium. Interestingly, the substrate binding capacity of *E. tenella* M1Pase (*K_m_* = 0.07 mM) was lower than that of SjaM1Pase2 (*K_m_* = 0.02 mM), and its catalytic efficiency (k_cat_ = 430 s^-1^) was much lower than that of SjaM1Pase1 (k_cat_ = 6453.5 s^-1^), which indicated that the *M1Pase* genes from brown algae may be better candidates for microbial production. Furthermore, mannitol biosynthesis is one of the most extensively tested targets for improving salt tolerance in plants by genetic engineering (Iwamoto and Shiraiwa, 2005). The high salinity tolerance of transgenic plants may due to the accumulation of mannitol in the cells (Tarczynski *et al*., 1992, 1993; Karakas *et al*., 1997). The introduction of algal genes in transgenic plants may confer a greater advantage in terms of salt tolerance (Iwamoto *et al*., 2001). The analysis of genes in the mannitol synthesis pathway can provide enzymes with higher substrate specificity and specific activity that would be useful for plant breeding research in the future.

## Acknowledgements

This work was supported by the National Natural Science Foundation of China (NSFC No. 41376143), Leading Talents Program in Taishan Industry of Shandong Province, Leading Talents Program in Entrepreneurship and Innovation of Qingdao, China-ASEAN Maritime Cooperation Fund “China-ASEAN Center for Joint Research and Promotion of Marine Aquaculture Technology”, and China Agriculture Research System (CARS-50).

